# Steroid hormone agonists reduce female fitness in insecticide-resistant *Anopheles* populations

**DOI:** 10.1101/2020.02.14.949024

**Authors:** Faye Brown, Douglas G Paton, Flaminia Catteruccia, Hilary Ranson, Victoria A Ingham

## Abstract

Insecticide based vector control tools such as insecticide treated bednets and indoor residual spraying represent the cornerstones of malaria control programs. Resistance to chemistries used in these programs is now widespread and represents a significant threat to the gains seen in reducing malaria-related morbidity and mortality. Recently, disruption of the 20-hydroxyecdysone steroid hormone pathway was shown to reduce *Plasmodium* development time and significantly reduce both longevity and egg production in a laboratory susceptible *Anopheles gambiae* population. Here, we demonstrate that disruption of this pathway by application of methoxyfenozide (MET) to insecticide resistant *An. coluzzii, An. gambiae sl* and *An. funestus* populations significantly reduces egg production in both topical and tarsal application. Moreover, MET reduces adult longevity when applied topically, and tarsally after blood feeding. As the cytochrome p450s elevated in pyrethroid resistant *Anopheles* only bind MET very weakly, this compound is unlikely to be subject to cross-resistance in a field-based setting. Manipulation of this hormonal signalling pathway therefore represents a potential complementary approach to current malaria control strategies, particularly in areas where high levels of insecticide resistance are compromising existing tools.

## 1. Introduction

The most effective form of malaria prevention is the control of the insect vector, the *Anopheles* mosquito (Bhatt et al., 2015). Vector control relies heavily on pyrethroid insecticides, which are currently used in all long-lasting insecticidal nets (LLINs), the main malaria prevention tool. LLINs provide personal protection through barrier action, preventing a host-seeking female from contacting the bed net user (Hawley et al., 2003). They also reduce the longevity of adult females and the likelihood they will live long enough to acquire and transmit the *Plasmodium* parasite. The heavy use of pyrethroid insecticides in LLINs, in addition to their use for control of agricultural pests, has inevitably led to widespread resistance to this insecticide class; indeed, pyrethroid resistance is observed in malaria vectors in over 80% of all surveillance sites in the WHO African region (“World Malaria Report.,” 2019) and is likely reducing the effectiveness of LLINs (Protopopoff et al., 2018; Toé et al., 2014). There is a clear and urgent need for new chemistries with distinct modes of action that can be incorporated into LLINs in order to combat increasing insecticide resistance in natural populations.

Resistance to pyrethroid insecticides is caused by at least four characterised mechanisms, several of which can be present in an individual mosquito; changes to the insecticide target site (Martinez-Torres et al., 1998; Ranson et al., 2000), reduced penetrance due to thickening of the insect cuticle (Balabanidou et al., 2016); sequestration by chemosensory proteins (Ingham et al., 2019) and increased metabolic clearance by detoxification enzymes such as cytochrome p450s (Müller et al., 2008; Stevenson et al., 2011). To restore the efficacy of pyrethroid treated bed nets in areas with resistant vectors, next generation nets impregnated with both pyrethroid insecticides and the cytochrome p450 inhibitor piperonyl butoxide (PBO) (Protopopoff et al., 2018) are now being implemented in several African countries. In addition, dual action nets with pyrethroids plus the sterilising compound pyriproxyfen have been successfully trialled in a field settings (Bayili et al., 2017; Protopopoff et al., 2018; Tiono et al., 2018) and this class of nets are expected to be in field use by the end of 2020. Mathematical models demonstrate that these new classes of nets could play a key role in reducing malaria cases in areas with high levels of insecticide resistance (Churcher et al., 2016). Nevertheless, there is a continual need for innovation into new LLINs, to pre-empt future resistance challenges; indeed there are already reports of cross resistance between pyrethroids and pyriproxyfen which may undermine the performance of these dual action nets in some settings (Yunta et al., 2016).

Recently, a 20-hydroxyecdysone (20E) agonist, methoxyfenozide (MET), was shown to impact multiple aspects of *Anopheles* biology that are relevant to its ability to carry and transmit *Plasmodium* (Childs et al., 2016). 20E is a major steroid hormone that, in multiple *Anopheles* species, is important for the reproductive fitness of adult male mosquitoes (Pondeville et al., 2008), alongside its conserved roles in adult female oogenesis (Baldini et al., 2013; Klowden and Russell, 2004) and larval development (Gilbert and Rewitz, 2009). Natural male 20E transfer during mating triggers substantial molecular responses within the female (Pondeville et al., 2008) leading to dramatic physiological changes including a loss of mating susceptibility, immune modulation, disinhibition of oviposition behaviour (Klowden and Russell, 2004) and upregulation of genes associated with oogenesis (Baldini et al., 2013). In a previous study, direct, topical application of MET onto *An. gambiae* females (G3 strain) induced a multimodal cascade of effects comprising substantially reduced lifespan and fecundity, reduced mating susceptibility, and partial blockage of *Plasmodium falciparum* transmission (Childs et al., 2016). Mathematical modelling predicted that, due to the combined effect on multiple traits, the use of MET as a vector control tool could produce comparable reductions in malaria burden to insecticide-based vector control (Childs et al., 2016). However, to be effective in the field, proposed new chemistries for incorporation into LLINs must overcome two key challenges. Firstly, they need to demonstrate efficacy against multiple wild insecticide resistant populations. Secondly, as tarsal uptake is the major route of delivery of insecticides on LLINs, the insecticides must be able to traverse the insect cuticle to reach their target sites.

Here we evaluate the effects of MET treatment in a number of insecticide-resistant *Anopheles* populations from sub-Saharan Africa, focusing on two important female life history traits: oogenesis and lifespan (Lees et al., 2019). We show that, similar to previous results in insecticide-susceptible *An. gambiae*, topical application of MET significantly reduces the number of eggs and the adult lifespan of highly pyrethroid resistant populations. Importantly, similar effects were observed when using tarsal uptake bioassays in the presence of an adjuvant. These data provide evidence that chemistries manipulating 20E function could be viable additions to the toolbox for malaria control.

## 2. Materials and methods

### 2.1. Mosquito rearing

All tests were carried out using 2-5 day old non-blood fed female mosquitoes of the resistant *An. gambiae* s.l. strain Tiassalé (Edi et al., 2012) from Cote D’Ivoire, two *An. coluzzii* strains, VK72014 and Banfora, from Burkina Faso (Namountougou et al., 2012) and *An. funestus* FuMoz from Mozambique (Hunt et al., 2005). Details of the resistance levels and underpinning mechanisms in these strains are provided in (Williams et al., 2019). The lab susceptible *An. gambiae* G3 strain was used as a control in all experiments. Insectaries were maintained under standard insectary conditions at 26°C ± 2°C and 70% relative humidity ± 10% under L12:D12 hour light:dark photoperiod. All stages of larvae were fed on ground fish food (Tetramin tropical flakes, Tetra, Blacksburg, VA, USA) and adults were provided with 10% sucrose solution ad libitum.

### 2.2. Topical application or methoxyfenozide

An 8% stock solution of the compound methoxyfenozide (MET) (Sigma-Aldrich) was prepared in DMSO; this stock was diluted in acetone to a 0.4% to match the concentration used in previous studies (Childs et al., 2016). Mosquitoes were anesthetized using carbon dioxide, placed on a petri dish lined with filter paper with the thorax exposed and held on a 4°C chill table (BioQuip products, Rancho Dominguez, CA). Using a hand-operated micro applicator (Burkhard Scientific, Uxbridge, UK), 0.5-µl of the 0.4% MET solution or a solvent-only negative control was applied to the dorsal thorax using a 1 cm^3^ syringe. Mosquitoes were transferred into holding cups, supplied with 10% sucrose solution and held in insectary conditions. Mortality was scored 24 hours post topical treatment and daily thereafter.

### 2.3. Tarsal Contact Assay

A 0.1438% solution of MET was prepared in acetone alone or with 0.0392% RME. A 500μl aliquot of the insecticide solution or solvent-only control was applied to the surface of Petri dishes (radius 2.5cm, area 19.635cm^2^) resulting in a MET concentration of 367mg/m^2^ and where applicable, 100mg/m^2^ RME. The petri dishes were placed onto an orbital shaker for 4 hours to evenly coat the surface and allow the dish to dry. Treated surfaces were used for one day and discarded. A 25ml plastic pot with a hole cut through the base was inverted and used to cover the surface to contain the mosquitoes. Replicates of ten mosquitoes were introduced through the hole and exposed to the surface for 30 minutes in insectary conditions. Mosquitoes were aspirated from the surface and transferred to holding cups, maintained on 10% sucrose solution and mortality scored 24 hours after exposure. Mosquitoes were then used to test the effect on egg production and survival as for the topical assays.

### 2.4. Egg Counts

To test the effect of MET on egg production, 24-hour post exposure, surviving mosquitoes were fed on human blood using a Haemotek membrane feeding system. Those that unsuccessfully fed or were partially fed were removed from the experiment. 72 hours post-blood feeding, mosquitoes were knocked down in the freezer and then dissected on glass sides. A few drops of distilled water were used to aid the removal of the ovaries under a dissecting microscope. Ovaries were gently teased apart and the number of eggs were counted and scored. Egg counts were assessed using GraphPad Prism 7.0 software, raw counts were compared using a Mann-Whitney U test and the egg intensity compared using a chi-squared test.

### 2.5. Longevity

To assess survival post MET treatment, mosquitoes were transferred into cups of ten and supplied with sugar water. Mortality was recorded every 24 hours, with dead mosquitoes being removed daily and assays continuing until all mosquitoes had died. In separate experiments on G3, Tiassalé and Fumoz strains, mosquitoes were blood fed 24 hours post exposure to the treated surface and scored every 24 hours until all mosquitoes had died. Longevity was assessed using GraphPad Prism 7.0 using a Mantel-Cox test.

### 2.6. Cytochrome P450 competitive binding assays

Competitive binding assays with MET and the fluorescent molecule diethoxyfluorescein (DEF) were carried out as previously described (Yunta et al., 2019) using the identical enzyme batch used in published studies evaluating binding of other public health insecticides (Yunta et al., 2019). Methoxyfenozide concentrations ranged from 0.0013μM to 2000μM final concentration; control wells contained DEF only. Reactions were set up in triplicate in a 96-well flat-bottomed, black plate and fluorescent read out performed on a fluorimeter. IC_50_ values were calculated on GraphPad Prism 7.

## 3. Results

### 3.1. Topical application of methoxyfenozide disrupts egg production and decreases lifespan in multi-resistant *Anopheles* populations

Methoxyfenozide was applied topically to four insecticide-resistant *Anopheles* populations, alongside a susceptible control (*An. gambiae* G3). The four resistant populations included one *An. funestus* (Fumoz), one *An. gambiae s*.*l*. (Tiassalé), and two *An. coluzzii* populations (Banfora and VK7). All populations are resistant to pyrethroids but the underpinning mechanisms differ (Williams et al., 2019). 24-hours post-application, the females were blood fed and 72-hours later ovaries were dissected and egg numbers scored. In each population, the total number of eggs in the MET treated group were significantly lower (p _Mann-Whitney_ < 0.0001) than in the respective acetone treated controls (Figure 1a) and in some cases treatment induced almost total suppression of egg development. Furthermore, the proportion of blood fed females that failed to develop eggs was significantly higher in the treated than the control group for all strains (p χ^2^ < 0.0001; df = 1) (Figure 1b). Adult lifespan was also significantly reduced by MET treatment in all populations (p _Mantel-Cox_ ≤ 0.001; df = 1), with the exception of *An. funestus*.

**Figure 1.**
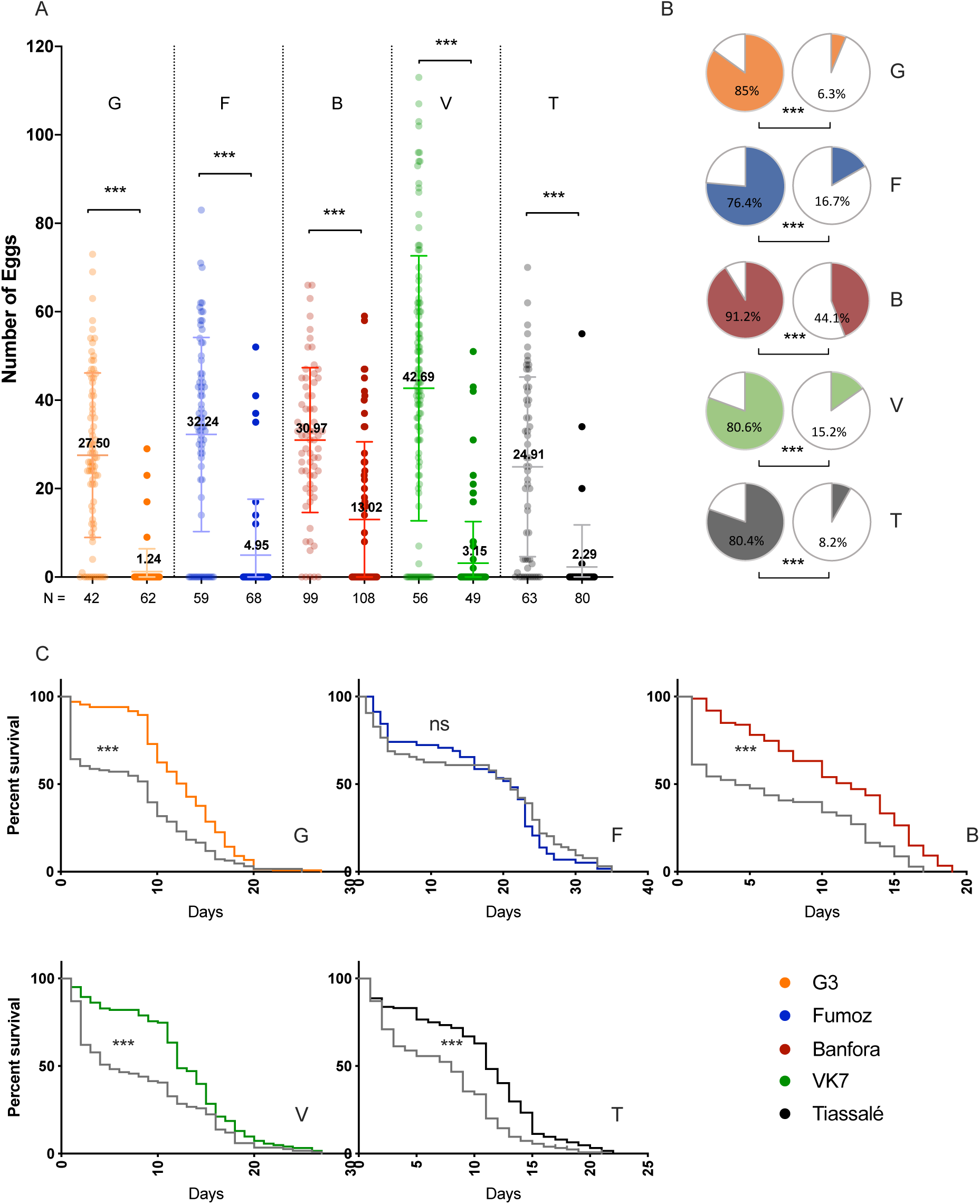
Topical application of 0.4% methoxyfenozide affects egg development and lifespan in multiple insecticide resistant *Anopheles* populations. A lab susceptible colony (G3: G); a pyrethroid-resistant *An. funestus* population (Fumoz: F); two pyrethroid-resistant *An. coluzzii* populations (Banfora: B and VK7: V); and a pyrethroid-resistant *An. gambiae sl* population (Tiassalé: T) were subjected to topical application of the 20E agonist 0.4% MET. A. MET-treated females (rightmost group) developed a significantly lower number of eggs than controls (leftmost group) in each population. *** p ≤ 0.001 as calculated by a Mann-Whitney U test, means and standard deviations are displayed for each group. Mean values are shown in each group. B. Treatment (right) resulted in a significantly lower number of females developing eggs in each population compared to controls (left). Percentage indicated in pie chart corresponds to percentage developing eggs and is represented by block colour; white shows no egg development. *** p ≤ 0.001 as calculated by χ^2^ test. C. All *An. gambiae sl* populations treated with topical DBH (grey) show a signficantly reduced lifespan compared to controls, *** p ≤ 0.001 as calculated by a Mantel-Cox test. There was no effect on *An. funestus* longevity.

### 3.2. Tarsal contact with MET induces significant reductions in oogenesis and lifespan in insecticide-resistant *Anopheles* populations

We next tested whether MET was also active via tarsal contact, using the susceptible G3 and the resistant Tiassalé and Fumoz populations. While tarsal exposure to MET alone did not significantly reduce oogenesis in these populations, we detected an effect on the lab susceptible G3 population on both total number of eggs (p _Mann-Whitney_ = 0.0410) and number of females developing no eggs (p χ^2^ = 0.0079; df =1) (Supplementary Figure 1). In an attempt to overcome barriers to penetration, tarsal bioassays were repeated in the presence of rapeseed methyl ester (RME), an adjuvant which prevents crystal formation on the glass surface, and improves compound uptake through the insect tarsi (Lees et al., 2019). With the addition of RME, there was significant reduction in the number of eggs in all three populations treated with MET (p _G3;Mann-Whitney_ = 0.0051; p _Fumoz;Mann-Whitney_ = 0.011; p _Tiassalé;Mann-Whitney_ < 0.0001) (Figure 2a) and significantly more Tiassalé females had complete loss of egg production post-exposure (p χ^2^ = 0.0011; df =1) (Figure 2b). Mosquito lifespan was not significantly affected by MET exposure, with or without RME (Supplementary Figure 1c) if mosquitoes did not receive a blood meal. However, when the females were blood fed 24 hours post exposure, a significant reduction in life span was seen in *An. gambiae sl* Tiassalé (p _Mantel-Cox_ < 0.0001; df =1) and *An. funestus* populations (p _Mantel-Cox_ = 0.0457; df = 1) (Figure 2c).

**Figure 2.**
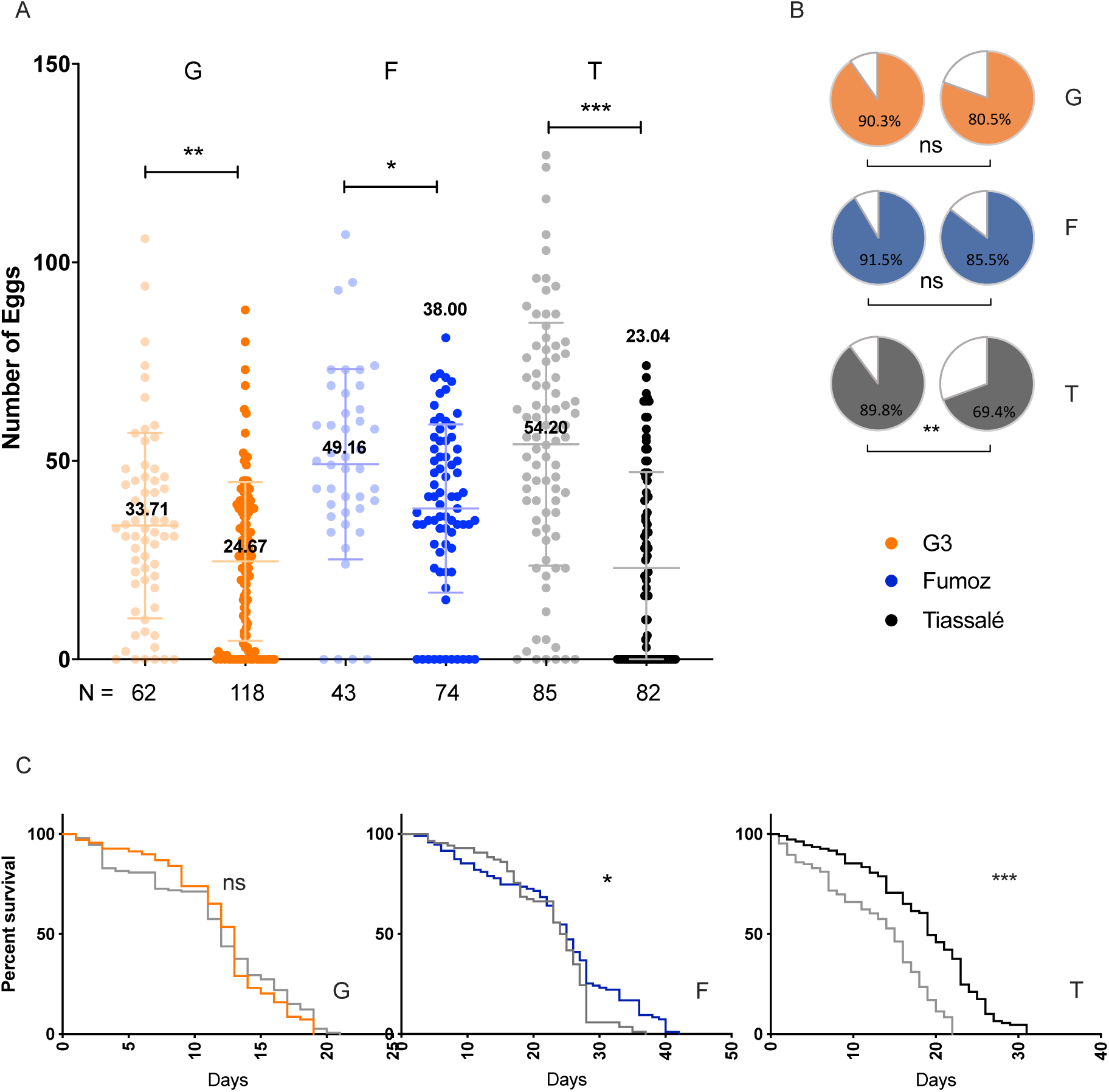
Disruption of steroid hormone signalling by tarsal application of methoxyfenozide + RME affects egg development and lifespan in multiple insecticide resistant *Anopheles* populations. A lab susceptible colony (G3: G); an *An. funestus* population (Fumoz: F); and an *An. gambiae sl* population (Tiassalé: T) were subjected to tarsal application of the 20E agonist methoxyfenozide, with RME. (A) Methoxyfenozide treated females (rightmost group) developed a significantly lower number of eggs than controls (leftmost group) in each population. * p ≤ 0.05; ** p ≤ 0.01; *** p ≤ 0.001 as calculated by a Mann-Whitney U test, means and standard deviations are displayed for each group. Means are shown for each group. (bB) Treatment (right) produced no significant changes in the number of females developing eggs in either the susceptible population nor *An. funestus*; however, the Tiassalé *An. gambiae s*.*l*. population showed a significant reduction in females developing eggs comapred to control (left). Percentage indicated in pie chart corresponds to percentage developing eggs and is represented by block colour; white shows no egg development. ** p ≤ 0.01 as calculated by χ^2^ test. (c) Bloodfed, treated resistant females (grey) show a signficantly reduced lifespan in both *An. gambiae s*.*l*. and *An. funestus* compared to controls, * p ≤ 0.05, *** p ≤ 0.001 as calculated by a Mantel-Cox test. there was no effect of methoxyfenozide on longevity in the susceptible *An. gambiae G3 strain*.

### 3.3. MET is not metabolised by the major detoxification enzymes cytochrome P450s

Cytochrome P450s are commonly upregulated in multiple resistant populations (Ingham et al., 2018) and are able to directly bind and metabolise insecticides including pyrethroids and pyriproxyfen (Yunta et al., 2016). To test whether the upregulation of these enzymes would inhibit MET function in a field setting, we performed competitive binding assays with six cytochrome P450s from the *An. gambiae* s.l. and one from *An. funestus* with the fluorescent substrate diethoxyfluorescein (DEF). The P450s tested include direct pyrethroid metabolisers *CYP6M2* (Stevenson et al., 2011), *CYP6P3* (Müller et al., 2008), and *CYP6P9a* (Ibrahim et al., 2016). All of the 7 cytochrome P450s tested showed weak binding affinity for MET (IC_50_ > 10 μM) (Table 1), with five of the P450s, including the *An funestus CYP6P9a* tested having an IC_50_ > 200μM and *CYP6P3* demonstrating no binding activity at all, suggesting that this chemistry is not readily metabolised by the P450s responsible for pyrethroid resistance.

## 4. Discussion

The identification of novel compounds to target the *Anopheles* mosquito is of utmost importance in order to maintain the gains in malaria control and accelerate progress towards elimination. Following on from earlier laboratory studies that demonstrated that topical application of the 20E agonist methoxyfenozide caused sterility and a reduction in lifespan in female *An. gambiae* mosquitoes (Childs et al., 2016), we demonstrate that this chemistry has a similar effect on multiple insecticide resistant *Anopheles* populations. Crucially, we also show that, with the addition of the adjuvant RME, methoxyfenozide interrupts egg development through tarsal contact Furthermore, blood fed females had a significant reduction in life span post methoxyfenozide exposure. Given that *Anopheles* mosquitoes are anautogenous, meaning they have to blood feed at least once to lay eggs, and need at least two blood meals to transmit pathogens, these findings of a small, but significant reduction in longevity may have important implications for vectorial capacity. The median survival time of Tiassalé and that of Fumoz fell in a lab-based setting; however, the reductions need to be confirmed under field conditions. A reduction in adult longevity is key to the community impact of LLINs as it reduces the probability that females survive for the duration of the parasite’s extrinsic incubation period in the mosquito. The direct impact on *Plasmodium* development was not investigated in the current study but previous work on the insecticide susceptible G3 strains found that methoxyfenozide exposure reduced the proportion of mosquitoes developing oocysts after feeding on a *P. falciparum* infected blood meal under laboratory settings (Childs et al., 2016). If this effect is also maintained in resistant mosquitoes under field settings, the use of methoxyfenozide would further decrease the likelihood of malaria transmission directly via an impact on the parasite and indirectly by shortening lifespan and reducing mosquito numbers in subsequent generations.

**Table 1.** Binding activity of insecticide resistance linked cytochrome P450s against methoxyfenozide. A range of MET concentrations ranging from 2000μM to 0.0013μM were used in competitive binding assays with the fluorescent substrate DEF with cytochrome p450s previously linked with insecticide resistance in two mosquito species, *An. gambiae* sl and *An. funestus* sl, in order to calculate IC_50_ values. Classification of binding strength is as follows: Strong inhibitors (IC_50_ < 1 μM); Moderate inhibitors (IC_50_ 1-10 μM); Weak inhibitors (IC_50_ > 10 μM). Gene identifier and 95% confidence interval of the IC_50_ values are also shown. IC50s of each p450 to pyriproxyfen is also shown, taken from (Yunta et al., 2019, 2016).

| Cytochrome P450 | Species | Identifier | IC<sub>50</sub> (μM) | 95% Confidence Interval | Pyriproxyfen IC<sub>50</sub> (μM) <sup>+</sup> |
| --- | --- | --- | --- | --- | --- |
| CYP6M2 | *Anopheles gambiae* sl | AGAP008212 | 15.75 | 10.12 to 24.38 | 14.14 |
| CYP9K1 | *Anopheles gambiae* sl | AGAP000818 | 26.39 | 16.60 to 41.46 | - |
| CYP6P4 | *Anopheles gambiae* sl | AGAP002867 | 251.4 | 103.7 to 657.6 | - |
| CYP6Z2 | *Anopheles gambiae* sl | AGAP008218 | 578.2 | 409.7 to 846.8 | 2.23 |
| CYP6P3 | *Anopheles gambiae* sl | AGAP002865 | No binding | No binding | 15.82 |
| CYP6Z3 | *Anopheles gambiae* sl | AGAP008217 | 208.7 | 99.05 to 475.1 | - |
| CYP6P9a | *Anopheles funestus* | AFUN015792 | 309.6 | 137.3 to 740.3 | 9.9 |

+ pyriproxyfen data collected as part of a different study, see <sup>16</sup>.

This study found that the use of the adjuvant RME was critical to detecting a physiological effect of MET via tarsal contact. RME acts as an emulsifier, reducing crystallisation of the chemical and aiding uptake (Lees et al., 2019). This adjuvant was included as a proof of principle to demonstrate that improving formulation could overcome barriers to uptake. Further work with alternative adjuvants and formulations would likely lead to marked improvements in uptake and effect sizes potentially similar to those observed with topical application.

All *An. gambiae* strains used in this study have multiple resistance mechanisms including target site and metabolic resistance, whereas in the *An. funestus* strain resistance appears to mediated solely by elevated cytochrome P450 activity (Williams et al., 2019). We observed a minimal interference from these insecticide resistance mechanisms on the effectiveness of MET, suggesting that a MET-based field-intervention could be effective in areas where conventional insecticides have reduced effectiveness. The juvenile hormone agonist, pyriproxyfen, is a more potent inhibitor of P450 enzymes (Yunta et al., 2019, 2016) and, direct metabolism of PPF by P450s has been demonstrated *in vitro* and associated with a moderate level of PPF resistance *in vivo* (Yunta et al., 2016). The results of the current study provide encouragement that the 20E agonist, MET, would be active against a wide range of field populations and may have reduced resistance liabilities compared to PPF. Whilst resistance to 20E analogs has been reported in agricultural pests (Ishtiaq et al., 2012; Mosallanejad and Smagghe, 2009; Rehan and Freed, 2014; Smagghe et al., 2012), the fitness cost is such that the effects are reversed soon after removal of this selection pressure (Rehan and Freed, 2014; Smagghe et al., 2012) which may enhance the durability of a 20E agonist-based intervention.

The public health value of incorporating insect growth regulators into LLINs has been previously demonstrated in a clinical trial of a PPF permethrin net from Sumitomo Chemical Ltd (Tiono et al., 2018). This trial, conducted in an area of intense malaria transmission and highly pyrethroid resistant vectors, found that malaria incidence was reduced in clusters using PPF-permethrin nets compared to standard LLINs. Further trials of PPF-permethrin nets are ongoing in Tanzania and Benin (London School of Hygiene and Tropical Medicine, 2018). However, given that potential for cross resistance between pyrethroids and PPF, incorporation of alternative insect growth regulators into LLINs should be considered as part of an insect resistance management strategy.

In conclusion, 20E agonists are a promising class of chemistry for use in vector control. They have low toxicity to non-target organisms (Nakagawa, 2005), are effective against pyrethroid resistant populations and can traverse the insect cuticle. In addition to a potential role as an alternative to PPF in LLINs, other possible applications targeting outdoor resting or feeding mosquitoes, such as attractive targeted sugar baits or push pull approaches should be explored.

## Acknowledgements

We would like to thank Rachel Davies, Sara Elg and Marion Morris for insectary support.

## Author contribution

F.B and V.A.I performed all experiments. D.G.P, F.C, H.R and V.A.I conceived the experimental design. All authors contributed to drafting of the manuscript.

---

**Extended Data Figure 1.**
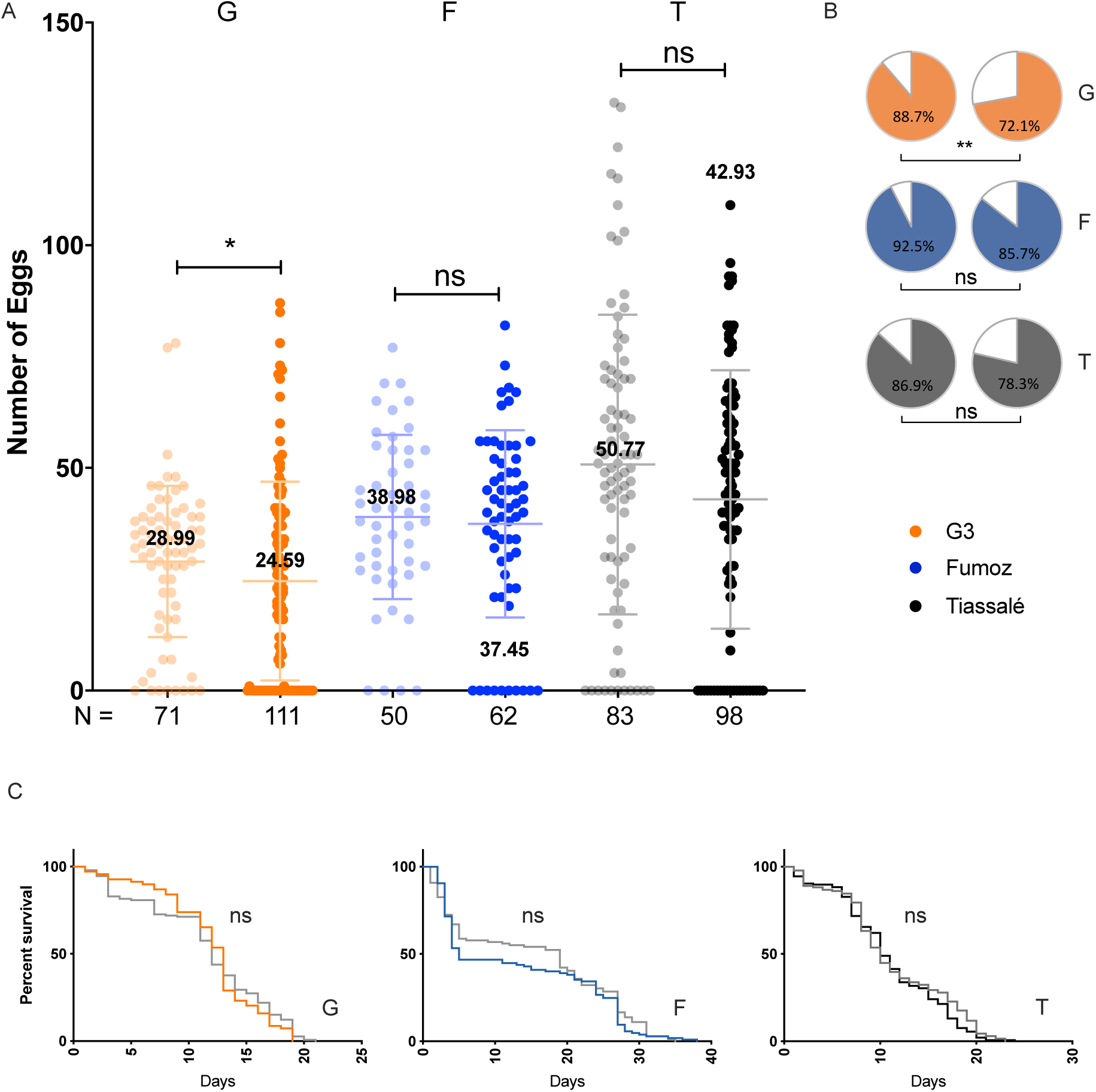
Tarsal exposure to 0.4% methoxyfenozide has no effect on multi-resistant *Anopheles* populations. A lab susceptible colony (G3: G); a multi-resistant *An. funestus* population (Fumoz: F); and a multi-resistant *An. gambiae sl* population (Tiassalé: T) were subjected to tarsal application of the 20E agonist methoxyfenozide (A) Methoxyfenozide treated females (rightmost group) developed a significantly lower number of eggs than controls (leftmost group) in the lab susceptible population G3 but no signficant effect was seen in either resistant colony. * p ≤ 0.05 as calculated by a Mann-Whitney U test, means and standard deviations are displayed for each group. (B) Treatment (right) produced no significant changes in the number of females developing eggs in either of the multi-resistant populations (left); however a signficant effect was seen on the lab susceptible population. ** p ≤ 0.01 as calculated by χ^2^ test. (C) *An. gambiae s*.*l*. and *An. funestus* females treated with tarsal DBH and RME (grey) show no change in lifespan compared to controls calculated by a Mantel-Cox test.

